# A roadblock-and-kill model explains the dynamical response to the DNA-targeting antibiotic ciprofloxacin

**DOI:** 10.1101/791145

**Authors:** Nikola Ojkic, Elin Lilja, Susana Direito, Angela Dawson, Rosalind J. Allen, Bartlomiej Waclaw

## Abstract

Fluoroquinolones - antibiotics that cause DNA damage by inhibiting DNA topoisomerases - are clinically important, but their mechanism of action is not yet fully understood. In particular, the dynamical response of bacterial cells to fluoroquinolone exposure has hardly been investigated, although the SOS response, triggered by DNA damage, is often thought to play a key role. Here we investigate growth inhibition of the bacterium *Escherichia coli* by the fluoroquinolone ciprofloxacin at low doses (up to 5x the minimum inhibitory concentration). We measure the long-term and short-term (dynamic) response of the growth rate and DNA production rate to ciprofloxacin, at both population- and single-cell level. We show that despite the molecular complexity of DNA metabolism, a simple `roadblock-and-kill’ model focusing on replication fork blockage and DNA damage by ciprofloxacin-poisoned DNA topoisomerase II (gyrase) quantitatively reproduces long-term growth rates. The model also predicts dynamical changes in DNA production rate in wild type *E. coli* and in an SOS-deficient mutant, following a step-up of ciprofloxacin. Our work reveals new insights into the dynamics of fluoroquinolone action, with important implications for predicting the rate of resistance evolution. Most importantly, our model explains why the response is delayed: it takes many doubling times to fragment the DNA sufficiently to inhibit gene expression. Our model also challenges the view that the SOS response plays a central role: the dynamical response is controlled by the timescale of DNA replication and gyrase binding/unbinding to the DNA rather than by the SOS response. More generally, our work highlights the importance of including biophysical processes in biochemical-systems models to fully understand bacterial response to antibiotics.

## Introduction

It is impossible to exaggerate the impact antibiotics have had on modern medicine, yet how exactly they inhibit bacteria remains controversial (Keren et al., 2013; Kohanski et al., 2007). Understanding the mechanism of antibiotic-induced growth inhibition is not only interesting from a basic science point of view, but also has the potential to contribute to rational drug design and optimization of treatment strategies that reduce the chance of resistance evolution (Chung et al., 2006; Ena et al., 1998; Greulich et al., 2015; Lukačišinová and Bollenbach, 2017; Meredith et al., 2015; Redgrave et al., 2014; Tan et al., 2012). To this end, quantitative models for antibiotic action that can be integrated into models for resistance evolution are much needed.

Even though many antibiotics have well-defined molecular targets (Boolchandani et al., 2019), the transition from a healthy bacterial cell to a dead, or non-growing, cell upon exposure to an antibiotic can be a complex and slow process. A prominent example is the bacterial response to fluoroquinolones – a class of DNA-targeting antibiotics that are used to treat a wide range of bacterial infections (Finch et al., 2012). Fluoroquinolone antibiotics typically produce a delayed response: bacteria initially continue to elongate after exposure (Elliott et al., 1987), and a significant fraction of cells are still viable after 2-3h (Wickens et al., 2000), even at doses where the antibiotic eventually kills almost all cells. Such a delayed response may play a role in the evolution of resistance, because elongating cells can continue to mutate and produce resistant offspring (Bos et al., 2015). However, no model has yet been proposed that explains the delayed response, and the delay also has not been accounted for in models for resistance evolution.

Fluoroquinolones target bacterial topoisomerases II (gyrase) and IV: enzymes that cut and re-seal the DNA, releasing the mechanical stresses accumulated during transcription and DNA replication, and helping to separate replicated chromosomes (Drlica and Zhao, 1997). Different fluoroquinolones have different binding affinities to topoisomerases II and IV. For example, ciprofloxacin – one of the most used antibiotics worldwide – binds predominantly to DNA gyrase in wild-type *E. coli* and only much more weakly to topoisomerase IV (Marcusson et al., 2009).

Ciprofloxacin traps the gyrase on the DNA as a DNA-protein complex and prevents it from dissociating (Drlica et al., 2009). This has two main effects. Firstly, the poisoned (ciprofloxacin-bound) gyrases act as roadblocks for DNA replication forks (Wentzell and Maxwell, 2000), blocking DNA synthesis (Drlica et al., 2008) and causing double-strand DNA breaks (DSBs) via a “chicken-foot” mechanism (Michel et al., 2004). Secondly, the poisoned gyrases also cause double-strand DNA breaks independently of replication fork activity (Drlica et al., 2008; Zhao et al., 2006). A single unrepaired DSB can be lethal in *E. coli* (Cockram et al., 2015), but cells have mechanisms to repair DSBs. One of these is SOS-mediated repair via the RecBCD machinery (Baharoglu and Mazel, 2014). A side effect of the activation of SOS is the suppression of cell division. The resulting filament formation and a change of the typical aspect ratio from ≈ 4 (Ojkic et al., 2019) to > 10 is a characteristic signature of exposure to fluoroquinolones (Bos et al., 2015). Therefore it is often thought that the SOS response is central in understanding the action of fluoroquinolones. However, despite much work on the molecular mechanism of fluoroquinolone action, very little work has been done on the dynamics of growth inhibition when antibiotic-naïve cells are exposed to a fluoroquinolone, and as yet no models have been proposed to predict this dynamical response, despite its relevance for resistance evolution. Moreover, some molecular aspects of the response also remain unclear; in particular the relative importance of DNA replication, replication-dependent and replication-independent DSBs, and SOS-mediated DSB repair (Drlica et al., 2008).

Here we use a combination of experiments and computer simulations to show that key features of the action of ciprofloxacin on growing *E. coli* bacteria, including the delayed dynamical response, can be explained using a relatively simple model. Consistent with the molecular mechanism described above, our model posits that ciprofloxacin-poisoned gyrase causes DNA replication fork stalling, and both replication-dependent and -independent DSBs. Double stand breakage causes a decrease in the net DNA production rate. The model successfully reproduces the long-term response to ciprofloxacin (growth inhibition curve) and, crucially, also predicts the short-term dynamics of *E. coli* in response to ciprofloxacin upshift, on the population- and single-cell levels. Our model could be integrated into population-level models for resistance evolution; the response delay, during which mutations can arise, may well have significant effects on the rate of emergence of resistance. Interestingly, our model also challenges the view that the SOS response is central, suggesting instead that the SOS system, while important in setting the model parameters, does not determine the time scale of the response to ciprofloxacin.

## Results

### 1. Parabolic shape of the growth inhibition curve suggests a cooperative inhibition mechanism

To understand the response of *E. coli* to ciprofloxacin (CIP) we first measured the long-term (steady-state) growth rate at different CIP concentrations: the growth inhibition curve. Previous work (Regoes et al., 2004) indicated that the inhibition curve of *E. coli* could be modelled by a Hill curve with a plateau at low concentrations. However, these experiments might not have been in a state of balanced growth as the bacteria were exposed to CIP for only one hour.

To determine the steady-state growth rate for different CIP concentrations, we used two different methods (Figs. 1, S1). We first measured *E. coli* growth curves for a series of CIP concentrations by incubating bacteria in microplates (200 µl/well) in a plate reader, and sampling the optical density every few minutes over 1-2 days (Methods). We used two strains: the K-12 strain MG1655, and a mutant derivative AD30. AD30 does not produce functional fimbriae and therefore sticks less to surfaces (Fig. 1B and Methods), preventing biofilm growth during the experiment. To minimize potential problems such as the dependence of optical density on cell shape (Pilizota et al., 2016), which changes during CIP-induced filamentation (Bos et al., 2015; Nonejuie et al., 2013), we extracted growth rates from time shifts between growth curves for cultures with different initial cell density (Methods). Both strains produced very similar growth inhibition curves with a characteristic inverted-parabola-like shape (Fig. 1A, B). This shape is consistent with previous results for ciprofloxacin (Regoes et al., 2004) but differs from that produced by many other antibiotics (Greulich et al., 2015; Regoes et al., 2004).

**Figure 1.**
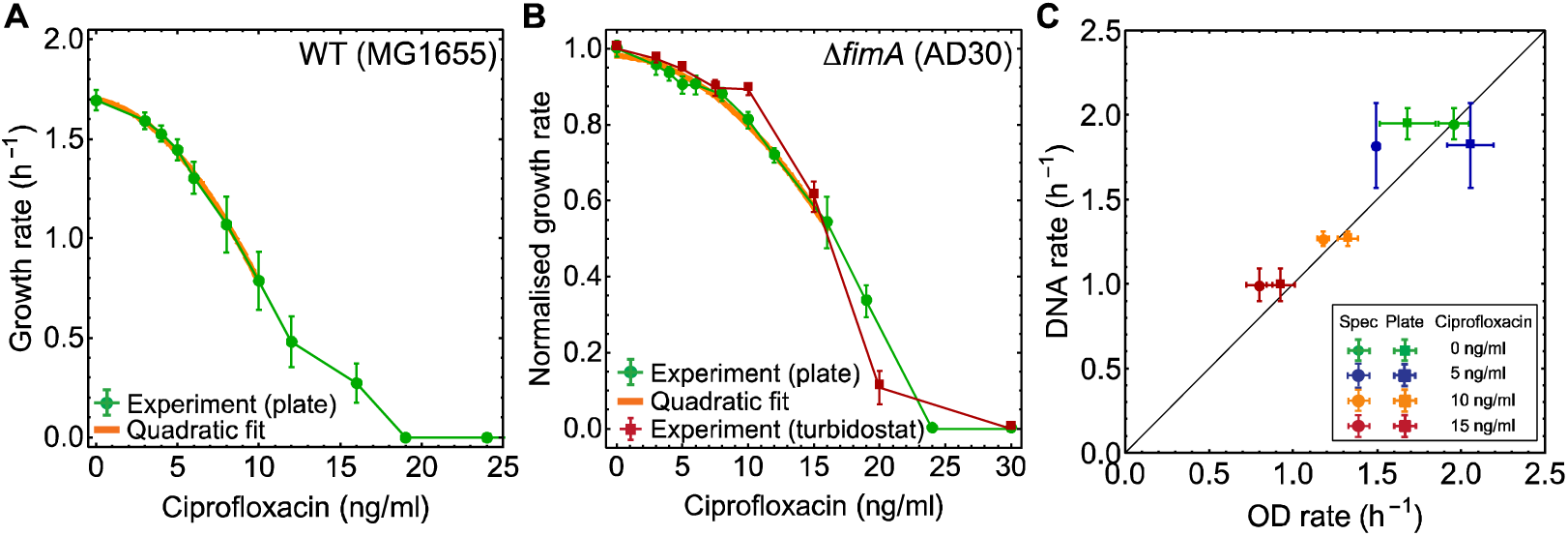
Growth-inhibition curve for ciprofloxacin and DNA production rates. (A) Growth-inhibition curve for ciprofloxacin treated *E. coli* (MG1655) for different antibiotic concentrations (plate reader data, green points). The orange line is a quadratic fit to the data. The minimum inhibitory concentration (MIC) is ∼20 ng/ml. Error bars represent SEM (4 replicates). (B) The growth-inhibition curve for the fimbrial knockout mutant (AD30). Growth rates are normalized (divided) by the growth rate in the absence of CIP. Green points are plate reader measurements, red points are measurements from turbidostat-incubated exponential cultures, taken ∼4 h after first exposure to ciprofloxacin. Both methods give similar results. Error bars are SEM (4 replicates). The MIC of AD30 is ∼25 ng/ml. (C) DNA production rate (measured by DAPI staining) correlates well with biomass growth rate (measured by OD). Error bars are SEM (3 replicates).

In parallel, we measured exponential growth rates for a range of CIP concentrations using steady-state cells grown in a turbidostat – a continuous culture device that dilutes cells once they reach a threshold density, maintaining exponential growth over long times (Methods and Fig. S1C, D). This could only be done for strain AD30, because the wild-type strain MG1655 rapidly forms a biofilm in the turbidostat. The growth rates in the turbidostat agree with those obtained from plate reader growth curves (Figure 1B).

If a culture is in a state of balanced exponential growth, all components of the bacterial cell must replicate at the same rate (Schaechter et al., 2006). Therefore the measured exponential growth rate should be the same as the rate of DNA synthesis. To confirm this, we measured total DNA at multiple time points in an exponentially growing culture for different CIP concentrations, and extracted the DNA production rate (Methods). Figure 1C shows that indeed the rate of DNA production matches the exponential growth rate as measured in our plate reader and turbidostat experiments.

Taken together, these results show that the long-term, steady-state rate of DNA production is a non-linear, inverted parabola-like function of CIP concentration, with only a small slope at zero CIP. If each DSB caused by CIP contributed (with probability) independently to the probability of cell death, and the number of DSBs was, the per-cell death rate would be proportional to. Assuming that increases proportionally to the CIP concentration, we would then expect a concave relationship between the net growth rate (birth minus death) and, with a negative slope at low. As this is not the case, a cooperative effect may be at play, which causes the number of DSBs to increase faster than linearly with. Alternatively, one might imagine a mechanism in which the number of DSBs is proportional to but must exceed a certain threshold before its effects on the growth rate become visible. We will show that the first hypothesis (non-linear increase of DSBs) is strongly supported by the data (Secs. 2-6), whereas the alternative hypothesis (threshold number of DSBs needed for growth inhibition) is not (Sec. 7).

### 2. A quantitative model for the action of ciprofloxacin

To understand how the rate of DNA synthesis is affected by ciprofloxacin, we developed a quantitative model (Fig. 2). The model includes reversible replication fork stalling by CIP-poisoned gyrases, and both replication-dependent and replication-independent double strand breakage.

**Figure 2.**
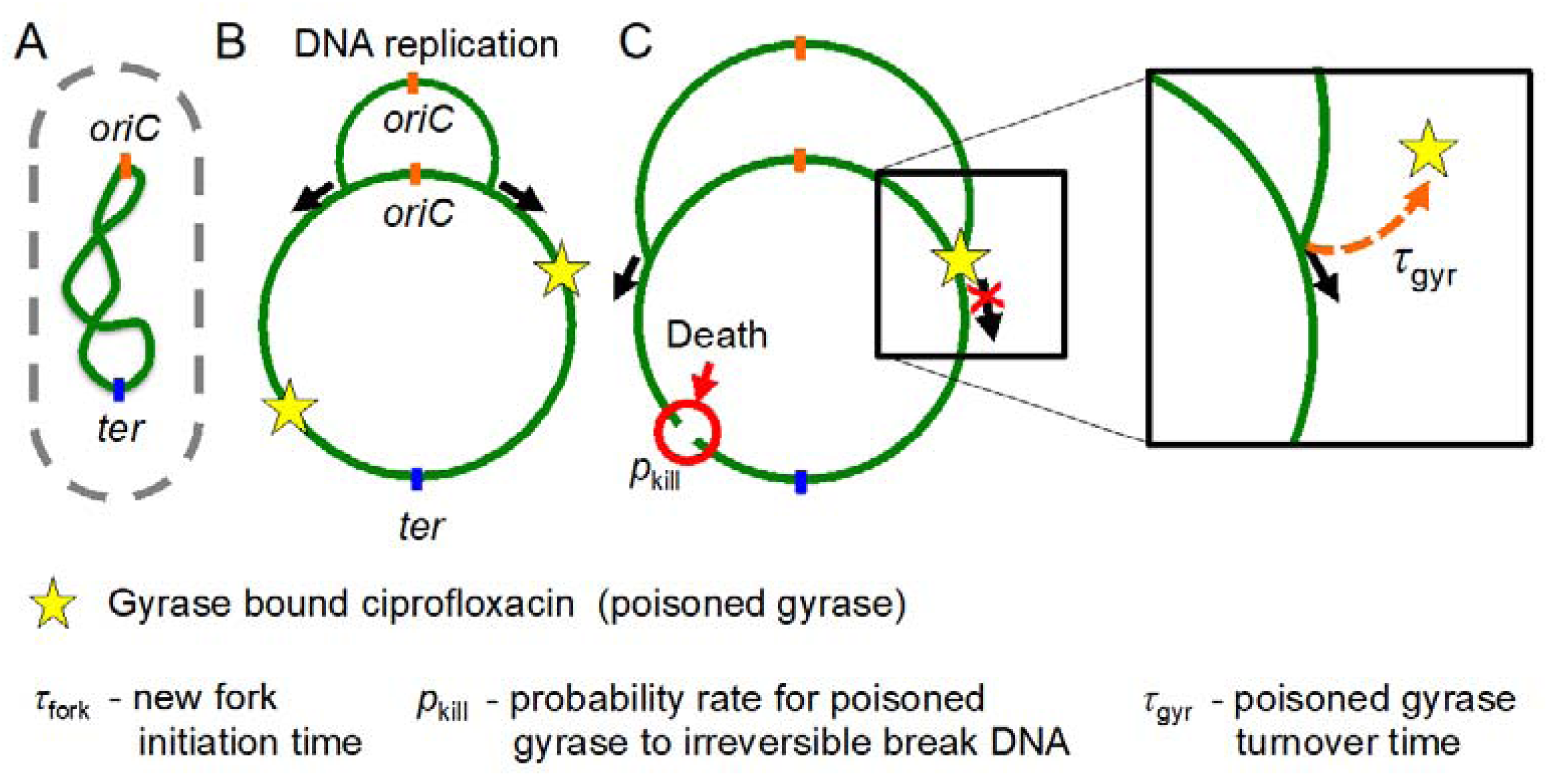
Model of ciprofloxacin mechanism of action. We model a collection of replicating chromosomes. New DNA is synthesized at replication forks (black arrows). Replication starts at the origin (*oriC*) and terminates at chromosome terminus (*ter*) (A). A newly synthesized DNA strand remains connected with the parent chromosome until the forks reach *ter* (B). Initiation of new forks at *oriC* occurs on average every *τ*_fork_ time units. The stars represent gyrases poisoned by ciprofloxacin. Poisoned gyrases are obstacles for replication forks, inducing fork stalling, and can also cause irreversible DNA damage with probability rate *p*_kill_ (C). Once poisoned gyrase is removed from the chromosome (with turnover time *τ*_gyr_), stalled forks resume replication.

In our model, a bacterial culture is represented by an ensemble of replicating circular chromosomes. New chromosomes are synthesized on the template of parent chromosomes and remain attached to them via replication forks. The forks start from the origin of replication (*oriC*) and end at the terminus (*ter*). Initiation occurs at time intervals drawn from a normal distribution with mean *τ*_fork_ =24 min chosen to reproduce the CIP-free growth rate from Fig. 1B, and standard deviation σ(*τ*_fork_)= 5 min (arbitrary value). Once initiated, replication forks progress at a constant rate *v*_f_ = 30 kb/min (Méchali, 2010). When a chromosome successfully completes replication, it separates from the parent chromosome.

Poisoned gyrases can appear anywhere along the chromosome with rate *k*_+_*L*/*L*_0_, where *k*_+_ is the DNA-poisoned gyrase binding rate, *L* is the current chromosome length, and *L*_0_ is the length of a fully replicated chromosome. We assume that the rate *k*_+_ is proportional to the extracellular CIP concentration *c* with an unknown proportionality constant *q* (units = 1/(time*concentration)): *k*_+_ = *qc*. Poisoned gyrases can also dissociate from the chromosome with rate *1*/*τ*_gyr_, where *τ*_gyr_ is the turnover time. The number of poisoned gyrases on the chromosome fluctuates, with the average value being determined by the balance between the binding and removal rates:

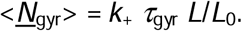

If a replication fork encounters a poisoned gyrase it stops and remains stalled until the poisoned gyrase is removed. The poisoned gyrase can also damage the entire chromosome irreversibly with rate *p*_kill_ (Fig. 2C). Damaged chromosome “conglomerates” (i.e. chromosomes plus any connected DNA loops) are removed from the simulation. The exact nature of the DNA damage is not important for the model, but a biologically plausible mechanism would be the creation of a DSB that does not get repaired (Drlica and Zhao, 1997). The process of repair is not modelled explicitly, but its effectiveness is implicitly included in the value of *p*_kill_ (e.g., a large value of *p*_kill_ corresponds to impaired DNA repair, since a poisoned gyrase is more likely to cause irreversible damage).

Our model has three unknown parameters: *τ*_gyr_, *p*_kill_, and the proportionality constant *q* that relates the extracellular concentration of CIP to the rate *k*_+_ with which poisoned gyrases appear on the chromosome.

### 3. The model reproduces the growth inhibition curve

We first checked if the model could reproduce the growth inhibition curve from Fig. 1. To do this, we calculated the rate of exponential increase of total DNA predicted by the model as a function of the CIP-proportional poisoned gyrase binding rate *k*_+_ (Fig. 3A, B). Figure 3B shows predicted growth inhibition curves for fixed *τ*_gyr_ = 15 min (arbitrary value) and a range of values of *p*_kill_. The simulated curves resemble the experimental curve (Fig. 1A). As expected, the rate of DNA synthesis decreases as the parameter *k*_+_ increases, mimicking increasing CIP concentration.

**Figure 3.**
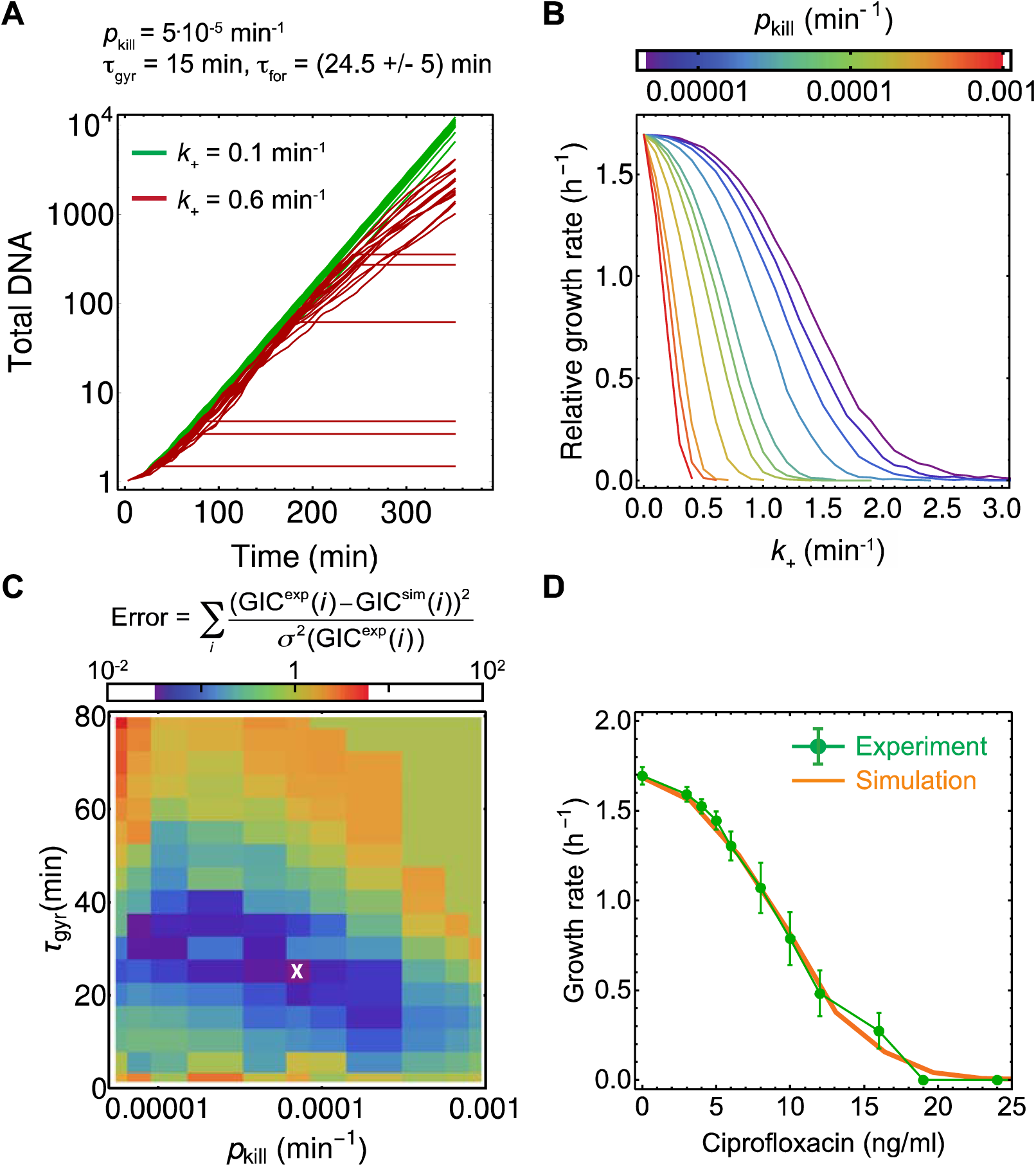
Simulations reproduce the measured growth inhibition curve. (A) Total amount of synthesized DNA predicted by the model as a function of time, for two different DNA-poisoned gyrase binding rates (*k*_+_ = 0.1 min^-1^, green, and *k*_+_ = 0.6 min^-1^, red). Total DNA is calculated as the total length of all chromosomes divided by *L*_0_. (B) Growth rate vs DNA-poisoned gyrase binding rate (*k*_+_) obtained by fitting exponential curves to the last 30 mins of the data from panel A, for different values of killing rate *p*_kill_. (C) Deviation between the experimental and modelled growth-inhibition curves as a function of *p*_kill_, *τ*_gyr_ (the third parameter, *q*, has also been fitted but is not shown). A cross marks the best-fit parameters *p*_kill_ = 7.10^−5^ min^-1^, *τ*_gyr_ = 25 min and *q* = 0.03 ml ng^-1^ min^-1^. (D) Experimentally measured growth inhibition curve (green), compared to the simulated best-fit curve (orange). Errors are SEM (four replicates).

We next systematically explored the parameter space (*p*_kill_, *τ*_gyr_, *q*) to find a range of parameter combinations that quantitatively reproduced our experimental data. Figure 3C shows that such a range indeed exists (dark blue region of Fig. 3C); the best-fit parameters are *p*_kill_ = (7 +/- 2).10^−5^ min^-1^ and *τ*_gyr_ = (25 +/- 5) min, and *q* = (0.030+/- 0.005) ml ng^-1^ min^-1^. This combination produces an excellent fit to the experimental data (Fig. 3D). Our fitted value for *τ*_gyr_ is about half the turnover time (∼55 min) that has been estimated from *in vitro* reconstitution assays (Kampranis and Maxwell, 1998); this discrepancy is perhaps not surprising since the *in vitro* assay lacks DNA repair systems (Baharoglu and Mazel, 2014) that may actively remove poisoned gyrases.

One can also extract from the model the average number of poisoned gyrases per chromosome, *N*_gyr_, for a given CIP concentration (Fig. S2). For a CIP concentration of 10 ng/ml, which corresponds to a two-fold reduction in the growth rate, we obtain *N*_gyr_≈4. The model thus suggests that a small number of poisoned gyrases is enough to inhibit growth.

Our model explains why the growth inhibition curve assumes a parabolic shape. At low concentrations of CIP there are very few poisoned gyrases present; DNA replication proceeds at almost normal speed and the chromosome topology is almost normal (since there are few blocked replication forks). Since the rate at which a chromosome conglomerate is damaged by CIP is proportional to the total DNA in the conglomerate, and *p*_kill_ is small, chromosome “death” is negligible at low CIP. However, as the CIP concentration *c* increases, replication forks become blocked more often. As a consequence, new replication forks are initiated before the parent and daughter chromosomes separate, producing large interconnected DNA conglomerates. Because the total DNA per conglomerate increases, the number of poisoned gyrases that are bound to the DNA also increases. This produces a faster-than linear increase in the degree of growth inhibition as *c* increases.

To confirm this interpretation of our model, we considered a modified model in which the damage caused by a poisoned gyrase does not “kill” the entire chromosome conglomerate but only the chromosome segment to which it is attached. There is some evidence that this might be the case for *E. coli* that is deficient in DSB repair (Sinha et al., 2018). This modified model predicts a very different growth inhibition curve (Fig. S3) which lacks the plateau at low CIP concentration.

### 4. The model predicts the dynamical response of *E. coli* to ciprofloxacin

Our model has been parameterized to reproduce the inhibition curve for steady-state growth in the presence of ciprofloxacin. To check if the model is able to predict the <u>dynamical</u> response of *E. coli* to ciprofloxacin (for which it has not been parameterized), we exposed strain AD30 to a step-up in ciprofloxacin concentration and measured dynamical changes in the growth rate over many generations in the turbidostat while maintaining cells in the exponential growth phase. Interestingly, we observed that for low concentrations of ciprofloxacin, the growth rate does not decrease immediately on antibiotic addition. The time until the growth rate begins to decrease, and the time to achieve a new steady-state growth rate, both depend on the CIP concentration (Fig. 4A).

**Figure 4.**
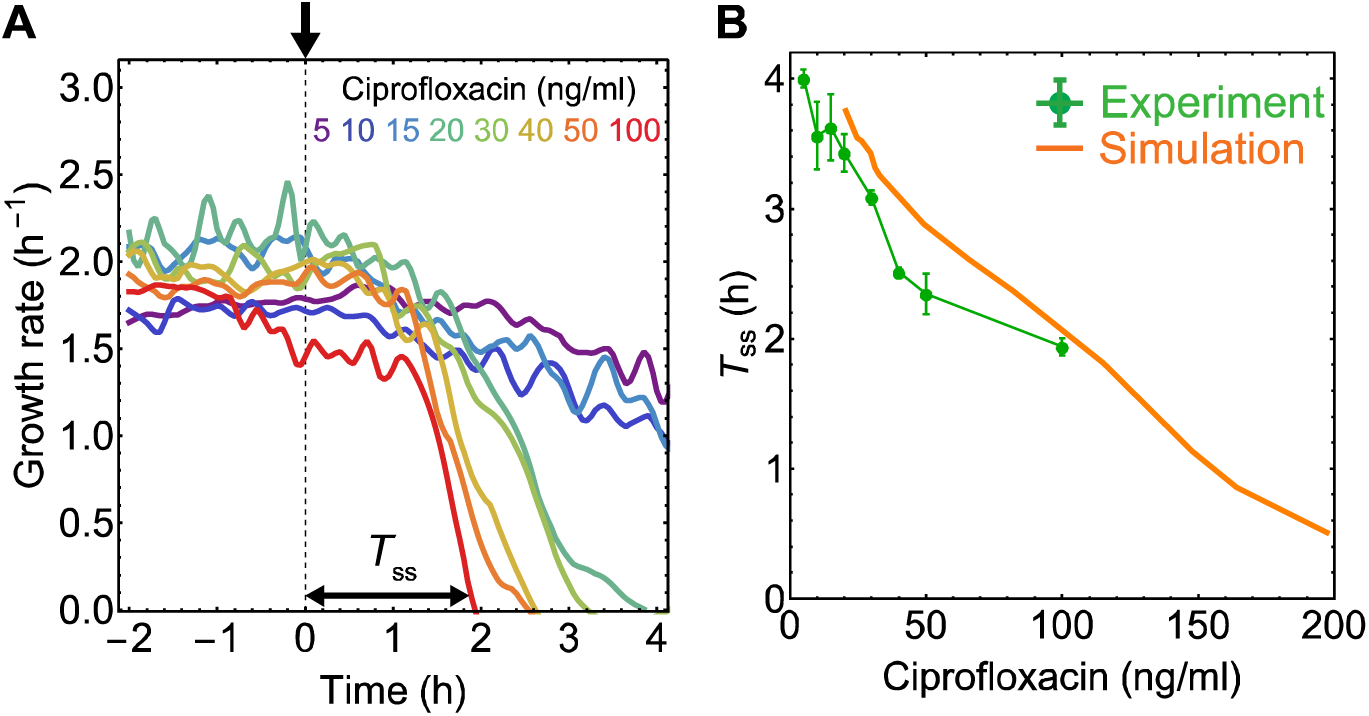
Dynamic response to CIP in the turbidostat. See Fig. S1C for a schematic diagram of the turbidostat. (A) Growth rate as a function of time for the fimbrial knockout strain AD30. Ciprofloxacin was added at 0 h. *T*_ss_ is the time to the new steady-state growth rate (*c* < MIC) or no growth (*c* > MIC) following the addition of CIP. (B) The model’s prediction for the time to new steady state is close to the experimental results.

Our model cannot predict the bacterial growth rate directly as it focuses on the rate of DNA synthesis, which does not have to be the same as the population-level growth rate during periods of unbalanced growth. However, the model can be used to predict the time to the new steady state. Figure 4B shows that the predicted values agree well with the results of our experiments, with no additional fitting parameters.

We next checked if the model also correctly predicts the dynamical response of DNA synthesis to ciprofloxacin exposure in single cells. We treated *E. coli* cells (MG1655) with ciprofloxacin for 1 hour, stained with DAPI to visualize DNA, and imaged in the bright field and fluorescent channels (Fig. 5). To prevent cell division and thus enable a direct comparison with the model, we used cephalexin (8 µg/ml), which inhibits PBP3, a component of the *E. coli* septation machinery (Pogliano et al., 1997). As expected, all the cells grew as filaments, without dividing (Fig. 5A).

**Figure 5.**
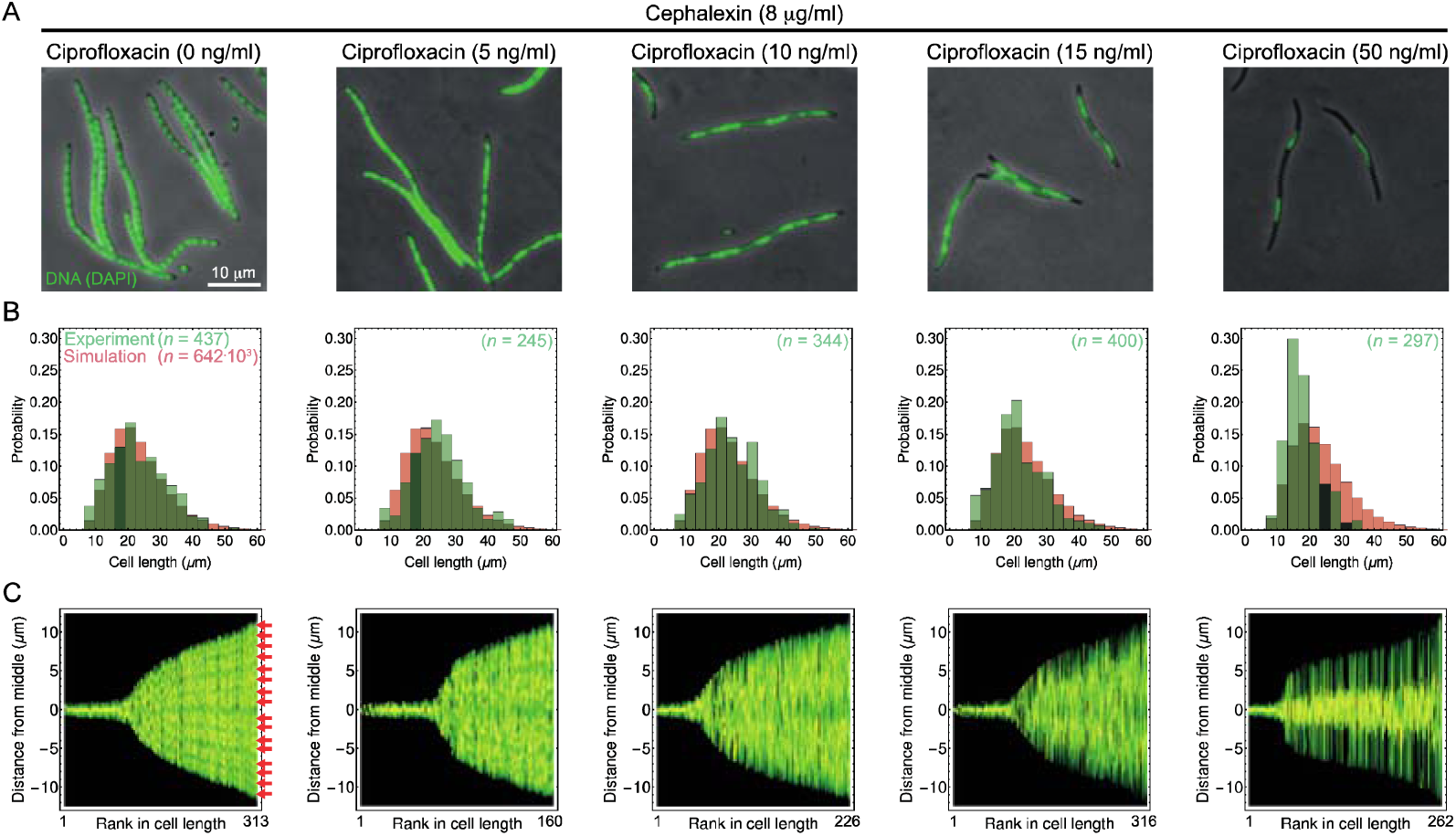
Ciprofloxacin causes formation of entangled DNA structures. (A) Phase contrast microscopy images overlaid with fluorescent DAPI stained DNA images, after 1 h exposure to different concentrations of ciprofloxacin. Cephalexin was added to prevent cell division (see Fig. S4 for CIP-only results). (B) Distribution of cell lengths after 1 h of CIP exposure (green = experiment, red = simulation). Cells shorter than 7 μm are excluded from the analysis. The best-fit for the cell length distribution for a CIP concentration of 50 ng/ml has *α* = 1.62 h^-1^, *σ*(*α*) = 0.07 h^-1^. Only the distribution for 50 ng/ml CIP differs from the CIP-free distribution (Kolmogorov-Smirnov *p*-value = 2.5.10^−15^ < 0.05). (C) Distribution of DNA in cells of different lengths. Cells are ordered by length from shortest to longest along the *x*-axis. Colour = DAPI fluorescence measured at different positions along cell midline (*y*-axis). Separate chromosomes (lighter areas pointed by red arrows) are clearly visible in CIP-untreated cells. The longest cells (∼24 μm) have ∼16 chromosomes. For 50 ng/ml CIP, chromosomes fail to separate (a single fluorescent region at cell’s midpoint).

The bacterial elongation rate is extracted from our measured filament length distributions by assuming exponential elongation with constant rate a starting from the initial length distribution of untreated cells (Methods). For cells treated with cephalexin only, the experimental length distribution was best fit by an elongation rate *α* =(1.85 +/- 0.28) h^-1^, similar to the growth rate obtained in plate reader experiments without any antibiotic (1.70 +/- 0.10 h^-1^, Figs. 1B, 5B). Therefore, cephalexin prevented cell division without visibly decreasing the biomass growth rate.

Remarkably, the cell length distribution (and hence the biomass growth rate) remained unchanged when cells were exposed to both ciprofloxacin (up to 15ng/ml) and cephalexin (Fig. 5B). This observation is consistent with previous microscopy data (Bos et al., 2015). Even at the highest CIP concentration used (50ng/ml, ∼2.5x MIC for this strain), the elongation rate was only slightly reduced (Fig. 5B, right).

We next characterized the DNA organization in single cells following exposure to CIP and cephalexin. Figure 5C shows that cells treated solely with cephalexin have clearly defined, evenly spaced chromosomes. The chromosome density is consistent with that of antibiotic-free growth; for example, for a cephalexin-treated filament of length 24 μm we observe ∼16 chromosomes, while *E. coli* of length 3 μm grown on LB antibiotic-free medium typically has ∼2 chromosomes (Fig. S5A). However, in the presence of CIP, DNA become less ordered and, as the CIP concentration increases, fewer distinct chromosomes can be identified. This suggests the presence of large entangled DNA structures and the failure of chromosome separation.

Our model makes a very specific prediction for how the total DNA in a filamentous cell should depend on CIP concentration after 1h of exposure (Fig. 6A). To test this prediction, we quantified the total DNA per cell by measuring DAPI fluorescence in microscopic images of cells for different concentrations of CIP. We obtained excellent quantitative agreement between our simulations and experiments (Fig. 6B), without any additional fitting. Thus our model, once fitted to the steady-state data, correctly predicts the early-time dynamical response to ciprofloxacin in single cells.

**Figure 6.**
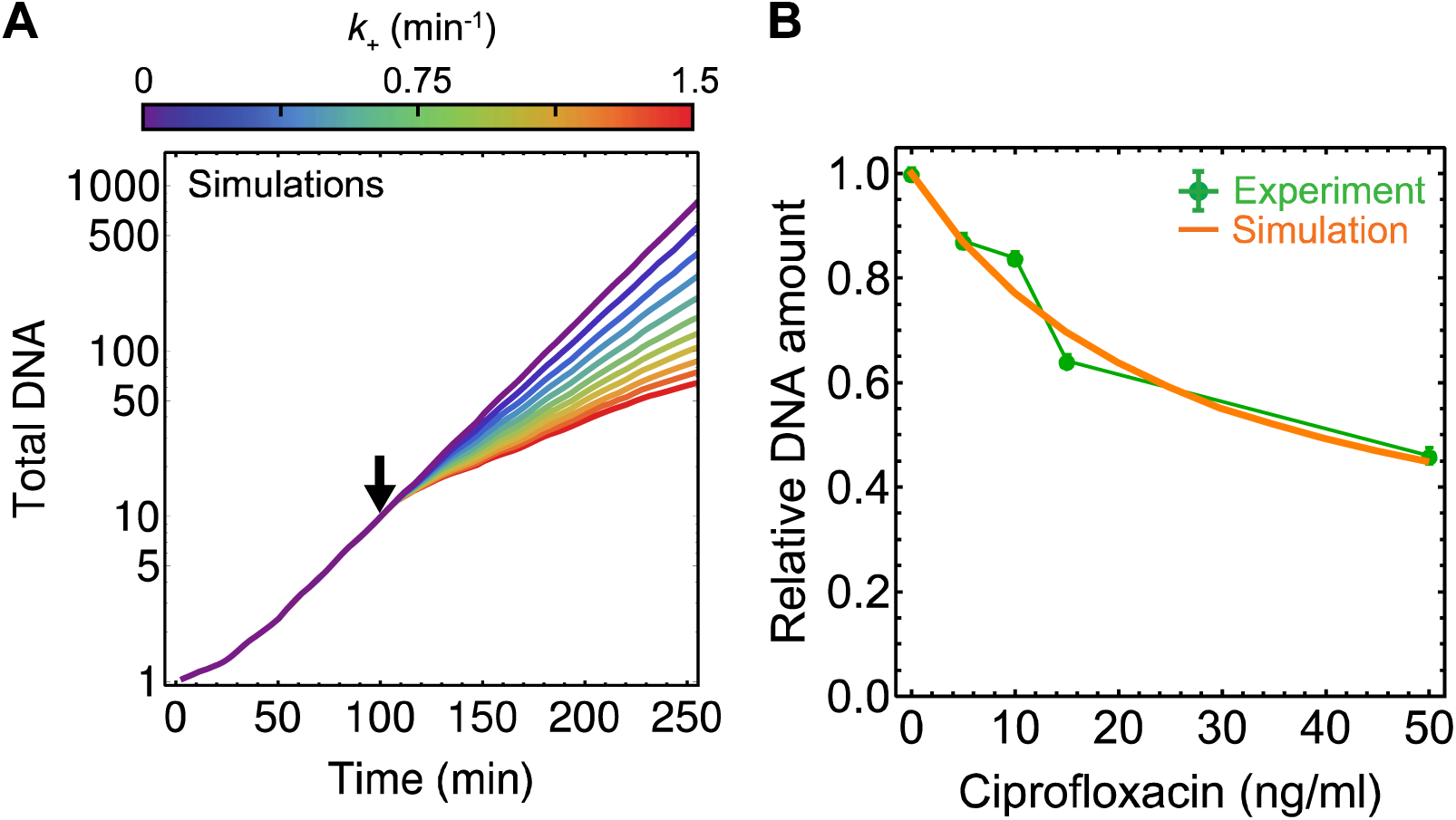
Simulations accurately predict the rate of DNA synthesis after ciprofloxacin exposure. (A) Simulated total DNA versus time (average of 1000 simulation runs). CIP is added at time *t* = 100 min. Different colors correspond to different gyrase binding rates k_+_ (different CIP concentrations). We used the best-fit parameters from Fig. 3. (B) Comparison of the predicted (no additional fitting) and experimentally measured total DNA per cell (DAPI staining) after 1 h of CIP exposure. Errors are SEM (350 cells on average per point).

### 5. Replication-dependent and replication-independent DNA damage predict the same shape of growth-inhibition curve

Ciprofloxacin-bound DNA gyrase has been hypothesized to cause both replication-dependent and replication-independent DNA double strand breaks (Drlica et al., 2008; Wentzell and Maxwell, 2000; Zhao et al., 2006). To test the role of replication-dependent versus replication-independent killing, we simulated a version of the model in which chromosome damage only occurs via fork-associated poisoned gyrase (Methods). This model turns out to reproduce the growth inhibition curve equally well (Fig. S6). Thus, replication-dependent or –independent DNA breaks produce the same growth inhibition dynamics.

### 6. Basal DNA damage is sufficient to model a DNA repair-deficient mutant

Our model does not explicitly include repair of DNA double strand breaks, which happens in *E. coli* via the RecBCD machinery, triggered by the SOS response (Courcelle and Hanawalt, 2003; Drlica and Zhao, 1997). We tested the role of DNA repair using a *recA* deletion mutant that cannot trigger the SOS response (Methods). We first investigated the growth of the Δ*recA* strain in the absence of ciprofloxacin. Δ*recA* cells were similar in length and width to WT cells, but had less organized chromosomes (Fig. S5B). In microplate cultures, the Δ*recA* strain showed a reduced growth rate compared to that of the WT MG1655 strain (∼1 h^-1^ versus 1.7 h^-1^ for WT). However, upon treating Δ*recA* cells with cephalexin and measuring the cell-length distribution after 1 h, we found that individual Δ*recA* cells elongate at the same rate as WT, although in the majority of the cells, the DNA looks less organized (Fig. 7A, B). To resolve this apparent contradiction, we imaged microcolonies of the Δ*recA* and WT strains growing on agar pads. Interestingly, the Δ*recA* colonies were significantly smaller and many colonies (∼30%) did not grow at all (Fig. S7). This suggests that the reduced population-level growth rate of Δ*recA* cultures is due to an increased fraction of non-growing cells, rather than a decreased single-cell growth rate. This is consistent with previous observations that cultures of bacteria deleted for *recA* can contain a significant sub-population of non-growing cells (Capaldo et al., 1974; Haefner, 1968).

**Figure 7.**
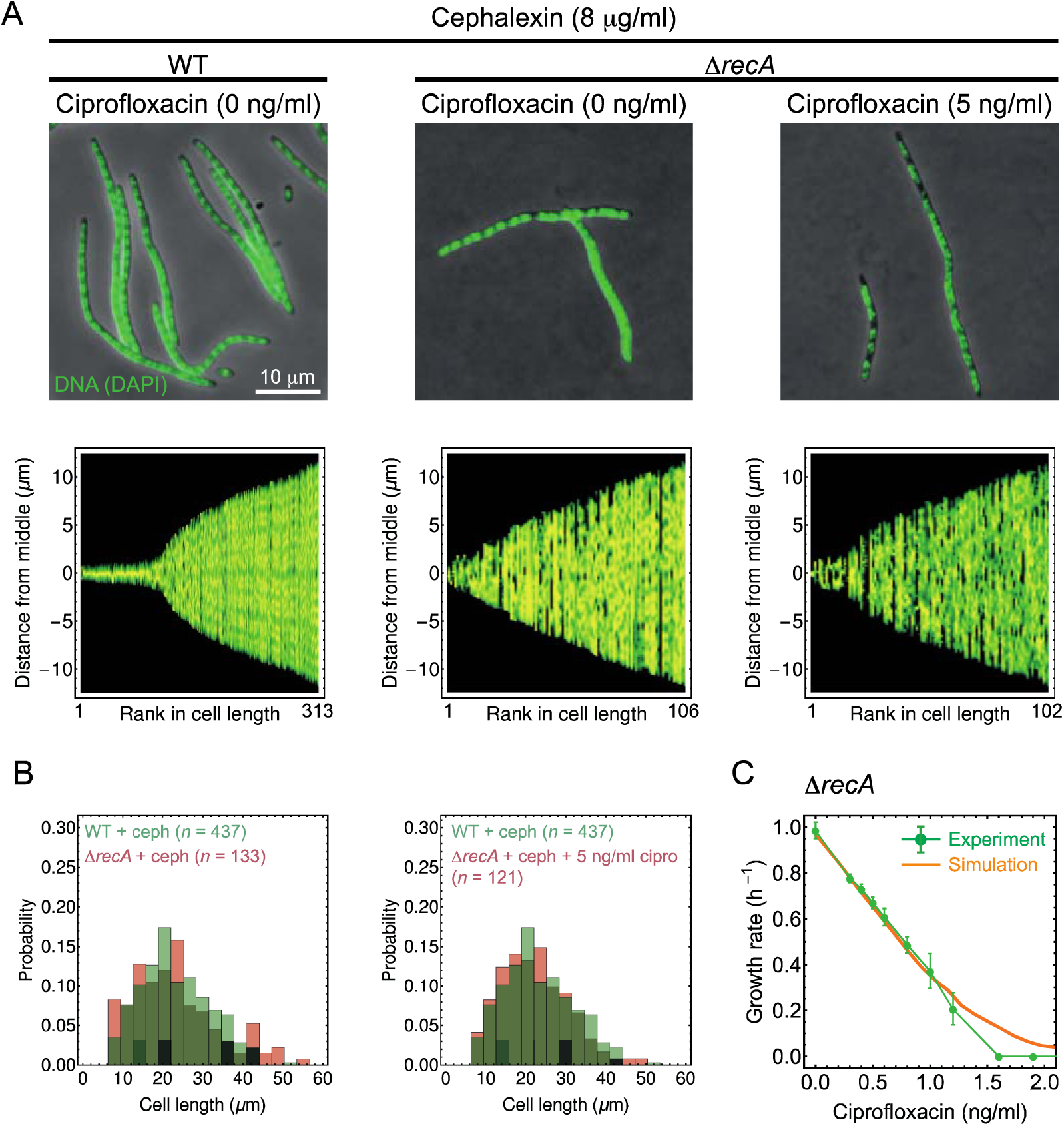
DNA-repair deficient cells (Δ*recA*) fail to separate chromosomes and are highly susceptible to ciprofloxacin. (A) Phase contrast microscopy images overlaid with fluorescent DAPI stained DNA images. All cells were treated with 8 μg/ml of cephalexin to prevent cell division. Many Δ*recA* cells fail to form separate chromosomes. (B) The cell-length distributions for Δ*recA* and WT after 1h of exposure to CIP do not differ even for a CIP concentration that inhibits the growth of Δ*recA* at the population level. (C) The model reproduces the experimental growth inhibition curve for Δ*recA.* Parameters *p*_kill0_ = (0.0033 +/- 0.0002) min^-1^, *p*_kill_ = (0.0042 +/- 0.0001) min^-1^ and *q* = 0.03 ml ng^-1^ min^-1^. Errors are SEM.

We next measured the growth inhibition curve for the Δ*recA* strain (Fig. 7C). The MIC of this strain (∼1.5ng/ml) was an order of magnitude lower than that of the WT. Moreover, the shape of the growth inhibition curve was significantly different than the WT parabola-like curve (Fig. 1): the growth rate decreased approximately linearly with increasing CIP concentration, without a plateau at low CIP. We hypothesized that these features could be reproduced in our model by an elevated rate of DNA damage associated with CIP-poisoned gyrases, combined with a basal DNA damage rate in the absence of CIP, both being due to the lack of the DSB repair mechanism. A modified model, in which the basal DNA damage rate *p*_kill0_ = 0.0033 +/- 0.0002 min^-1^ was fixed by fitting to the population growth rate in the absence of CIP, reproduced the experimental growth inhibition curve very well (Fig. 7C, *p*_kill_,*q* also fitted to the inhibition curve). This shows that even though our model does not explicitly include DNA repair, implicit dependence via the parameters *p*_kill0_ and *p*_kill_ is sufficient to reproduce our experimental data.

### 7. Alternative hypothesis based on saturation of repair mechanisms does not explain the data

Our model reproduces all our experimental observations – but could an alternative model based on a different microscopic mechanism explain them equally well? To investigate this, we considered a biologically plausible model in which the parabolic shape of the inhibition curve arises due to a non-linear response of the DNA repair mechanism to CIP concentration, rather than from a non-linearity in the amount of DNA damage as the previous model did.

In this alternative model, for CIP concentrations above the MIC, DSB repair mechanisms become saturated, which causes the accumulation of DSBs. Below the MIC, however, we assume that recBCD-mediated DSB repair (Michel and Leach, 2012) is very effective. Specifically, we assume that the number *n(t)* of DSBs evolves as

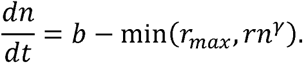

Here, DSBs are created at a rate *b* that is proportional to CIP concentration, and are removed via repair at a rate *rn*^*γ*^, which cannot exceed the maximum rate *r*_*max*_. The exponent *γ* characterizes the strength of the feedback between the number of DSBs and the rate of repair; *γ* = 1 corresponds to a linear response, whereas *γ* < 1 means that repair mechanisms are strongly triggered even by a small number of DSBs. We further assume that each DSB has equal probability *p* of killing the cell, hence the net growth rate is proportional to exp (−*pn*).

This model, which does not consider the dynamics of DNA replication, reproduces the steady-state growth inhibition curve quite well (Fig. S8) for *γ* ≈ 0.5. However, the model predicts that the time to reach a new steady-state growth rate following an upshift of CIP should be proportional to 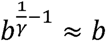. The time to the new steady state is thus predicted to increase with CIP concentration (since *b* increases with *c*) which disagrees with what we observe experimentally (Fig. 4). Therefore, this model fails to reproduce the dynamics of CIP inhibition.

## DISCUSSION

Despite much work on the molecular mechanisms of fluoroquinolone action, no models have yet been proposed to explain the delay in the bacterial response to low-dose exposure, even though this may well have important consequences for the chances of resistance evolution. We have proposed a quantitative model for fluoroquinolone-induced growth inhibition of the bacterium *E. coli* that for the first time explains the response delay. Our model is based on the known molecular details of replication fork stalling and DNA damage, and makes quantitative predictions for the long- and short-term (dynamic) bacterial response to the fluoroquinolone ciprofloxacin. By fitting the model’s three parameters (Fig. 3) to the experimental steady-state inhibition curve (long-term response), we not only reproduce the shape of the curve very well but we also make correct predictions for the short-term dynamics of bacterial growth following a step-up of ciprofloxacin. The predictions are in agreement with our experimental data, without any further parameter fitting (Fig. 4, 6, 7). Importantly, our model also challenges the view that the SOS DNA damage response plays a central role. Our model, with altered parameters, also reproduces the behavior of a *recA* mutant that cannot activate the DNA repair machinery and is significantly more sensitive to ciprofloxacin. Thus the SOS system can significantly alter the parameters of the model but, importantly, does not control the dynamics of the response. Instead, the dynamics is controlled by the DNA replication rate and binding/unbinding rates of gyrase from the DNA.

We have also considered modifications of the model to include DNA damage occurring only due to replication fork-associated gyrases, and damage that “kills” only the local DNA strand rather than the entire chromosome conglomerate. It turns out that our model cannot distinguish between fork-related and replication-independent killing, but is sensitive to whether poisoned gyrases kill the whole cluster of interconnected DNA, or only the local branch that is affected by a poisoned gyrase. The latter predicts a non-parabolic inhibition curve that is at odds with the experimental data. An alternative model based on the saturation of the repair mechanism as an explanation of the growth inhibition curve fails to predict the dynamic response to CIP.

Our work demonstrates that, despite the molecular complexity of fluoroquinolone action, a simple physiological model can explain the behavior of bacteria exposed to this class of antibiotics, leading to new insights that can be used to make quantitative predictions. Below we discuss in more detail some of the implications of our work.

### Shape of the growth inhibition curve

The growth inhibition curve for CIP is parabolic-like (Fig. 1). Inhibition curves for many antibiotics including CIP have been traditionally approximated using the Hill function (Regoes et al., 2004). This choice is often based on a qualitative resemblance of the shape rather than on mechanistic insight. The Hill function is also a popular choice in population-level models of antibiotic treatment (Levin and Udekwu, 2010; Lipsitch and Levin, 1997; Torella et al., 2010). However, some antibiotics can have very different inhibition curves, that are not well approximated by a Hill function (Greulich et al., 2015). This is potentially important for modelling the evolution of resistance to antibiotics, because differently shaped inhibition curves are expected to produce different fitness landscapes (Chevereau et al., 2015; Engelstädter, 2014), leading to different levels of selection for resistant mutants, and hence different trajectories to resistance.

We checked how well our measured growth inhibition curve can be reproduced using a Hill function (Fig. S9). The fit is slightly less good than that produced by our model. The Hill exponent (*κ* = 4.4 ± 0.5) also differs significantly from the value of *κ* = 1.1 ± 0.1 reported before (Regoes et al., 2004). We conclude that careful measurements of the steady-state growth inhibition curve, combined with physiological models of antibiotic response, can not only shed light on the mechanism of inhibition but are also required in quantitative models of the evolution of antibiotic resistance.

### The role of the SOS response

The cellular response to DNA damage is not explicitly included in our model, but rather enters through the parameter values. In others’ work, the SOS response has been modelled in the context of UV response (Aksenov et al., 1997; Belov et al., 2009; Krishna et al., 2007; Shimoni et al., 2009). To check how realistic it was to omit details of the SOS response in our model, we adapted the model from Ref. (Belov et al., 2009) to our scenario. We set the initial number of DSBs (parameter *N*_*G*_ from (Belov et al., 2009)) to zero, and added a term proportional to the CIP concentration to the equation *dN*_*G*_ /*dt* which describes the rate of change of the number of DSBs. We calculated the time it takes for LexA (the protein whose inactivation triggers the response) to reach a new steady state (10% above the infinite-time limit concentration). Figure S10 shows that this time is less than 10 mins for a broad range of DSB creation rates, indicating that the SOS response occurs much faster than the growth rate response we report in Fig. 4. Based on this and the excellent agreement between our model and experiments, we claim that key features of growth inhibition in response to sub-MIC ciprofloxacin (inhibition curve shape and inhibition dynamics) can be understood without modelling the SOS response explicitly. This does not mean that the SOS response is not important; on the contrary, SOS-induced changes in bacterial physiology (e.g., expression of low-fidelity polymerases) are very important for the evolution of resistance (Bos et al., 2015; Michel, 2005), and the role of SOS in mediating growth inhibition is also implicit in our model through the parameters *p*_kill_ and *p*_*kill*0_.

### The importance of chromosome segregation

In this work, we do not model individual cells; rather, we consider a collection of replicating chromosomes. While this seems to be enough to reproduce the population-level growth-rate response to ciprofloxacin, and the DNA dynamics in single cells, it cannot account for some aspects of behavior at the cellular level, such as the cell length distribution (in our experiments, we avoid this issue by treating cells with cephalexin). More work will be required to create a model that is able to, for example, predict the cell length distribution (Fig. 5), cell division and budding (Bos et al., 2015), or antibiotic-induced fluctuations in the number of cells in small populations (Coates et al., 2018).

### Relevance for bacterial infections

Predictive understanding of how antibiotics inhibit bacteria could help in the design of better treatment strategies. Traditionally, models for antibiotic treatment have assumed an instantaneous response of bacteria to the antibiotic (Bonhoeffer et al., 1997; Jumbe et al., 2003); models that take intracellular dynamics into account are still rare (Greulich et al., 2017; zur Wiesch et al., 2015). Our research shows that ignoring the transient behavior (here the short-term bacterial response delay) can be problematic because these transients can last for many generations at sub-MIC concentrations of the antibiotic for which the probability of developing resistance is the highest (Drlica, 2003; Greulich et al., 2017, 2012). Our physiological model could be integrated into population-level evolutionary models, allowing better prediction of the chances of resistance emergence that take account of the cell-level dynamical response. Such effects are almost universally neglected in current evolutionary models.

In conclusion, we have proposed and tested a model that predicts bacterial response to fluoroquinolones. Our model complements those that have recently been proposed for other classes of antibiotics; taken together, such models may eventually be useful in understanding and predicting bacterial response to clinically relevant treatment strategies, such as the effect of combination therapies (Bollenbach et al., 2009; Chait et al., 2007; Wood et al., 2012).

## STAR Methods

### Bacterial strains

We used MG1655, a K12 strain of the bacterium *E. coli*, and two mutant derivatives (AD30 and ΔrecA). Strain AD30 was constructed by P1 transduction from JW4277 (the *fimA* deletion strain in background BW25113 from the Keio collection) into MG1655 (Baba et al., 2006). The kanamycin resistance cassette was removed using Flp recombinase expressed in pCP20. Strain construction was confirmed by PCR using a combination of kanamycin specific primers and gene specific primers.

The Δ*recA* mutant was donated by M. El Karoui’s lab.: it is MG1655 in which ΔrecA::CmR was introduced by P1 transduction from DL0654 (David Leach, laboratory collection).

### Growth media and antibiotics

All our experiments were performed in LB medium at 37°C. LB liquid medium was prepared according to Miller’s formulation (10g tryptone, 5g yeast extract, 10g NaCl per litre). The pH was adjusted to 7.2 with NaOH before autoclaving at 121°C for 20 min. To create LB in 1.5% agar, agar (Oxoid, Agar Bacteriological, No. 1) was added before autoclaving.

Ciprofloxacin solutions were prepared from a frozen stock (10mg/ml CIP hydrochloride in ddH2O) by diluting into LB to achieve desired concentrations. Stock solution of cephalexin (10mg/ml) was prepared by dissolved 100mg of cephalexin monohydrate in 10 mL of DMSO.

### Growth inhibition curves

To determine the growth rate at a given concentration of CIP, we used two different methods.

#### Method 1

We incubated bacteria in a micro-plate inside a plate reader (BMG LABTECH FLUOstar Optima with a stacker) starting from two different initial cell densities, and measured the optical density (OD) of each culture every 2-5 min to obtain growth curves.

Plates were prepared automatically using a BMG LABTECH CLARIOstar plate reader equipped with two injectors to create different concentrations of CIP in each column of a 96 well plate (total injected volume 195μl per well). Bacteria were diluted from a thawed frozen stock 10^3^ and 10^4^ times in PBS, and 5μl of the suspension was added to each well (10^3^ dilution to rows A-D, 10^4^ dilution to rows E-H). After adding bacteria, plates were sealed with a transparent film to prevent evaporation, and put into a stacker (temperature 37°C, no shaking), from which they would be periodically fed into the FLUOstar Optima plate reader (37°C, orbital shaking at 200rpm for 10s prior to OD measurement).

Assuming that all cultures grow at the same rate when cell density is low (OD<0.1), the time shift (Δ*T*) between the curves from rows A-D and E-H (Fig. S1A) is related to the exponential growth rate as follows:

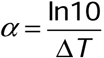

We used this relationship to calculate *α* from time shifts between 4 pairs of replicate experiments (A-E, B-F, C-G, D-H) for 12 concentrations of ciprofloxacin (range: 0—30 ng/ml). To validate the method we also calculated growth rates by fitting an exponential curve *A*+ *Be*^*αt*^ to the low-OD (OD<0.1) part of the growth curve. The time-shift method gives more accurate but overall similar results compared to the exponential curve fitting (Fig. S1B) or maximum growth rate measurement methods (Swain et al., 2016). Our fitting method is not sensitive to the relationship between the OD and the true cell density (which depends on the cell shape and size) and it gives the average growth rate over many more generations (growth from approx. 10^4^ to 10^8^ cells, ≈ 13 generations) than curve-fitting based methods (OD=0.01-0.1, 3 generations), see Fig. S1B.

#### Method 2

To confirm that our measurements correspond to steady-state growth, we also measured the growth rate in a turbidostat (Fig. S1C), in which bacteria are kept at approximately constant optical density (OD=0.075-0.1) for many generations by diluting the culture with fresh medium (and concomitant removal of spent medium + bacteria) whenever the OD reaches a threshold value. The growth rate is obtained by fitting an exponential function to the background-corrected OD data between consecutive dilutions.

We found that strains MG1655 and AD30 have similar but not identical susceptibility to ciprofloxacin: while the MG1655 wild type showed an MIC of (19 +/- 3) ng/ml, in agreement with previous measurements (Marcusson et al., 2009), AD30 was slightly less susceptible, with an MIC of (24+/- 3) ng/ml. The MIC values were determined from the zero-growth point of the growth inhibition curves (3-6 replicate experiments).

### Measurements of DNA production

To obtain the data in Fig. 1C, cells were grown in LB medium with or without CIP in shaken flasks (3 replicates), and diluted periodically with fresh medium to maintain steady-state exponential growth. Cells were sampled every ∼20-30 min, fixed (1ml of culture fixed with 250μL of 1.2% formaldehyde) and their OD was measured using both a standalone spectrophotometer (Cary 100 UV-Vis) and a plate reader (CLARIOstar) for cross-validation. DAPI was added to the fixed samples to a concentration 2 μg/mL (Nonejuie et al., 2013). After 30 min of incubation with DAPI the cells were washed 3 times with PBS, and DAPI fluorescence intensity was measured in the plate reader (CLARIOstar). Growth rates were extracted from the fluorescence and OD versus time curves by least-squares fitting of an exponential function.

### Microscopy

To obtain the data from Figs. 5 and 7, exponentially growing cells (LB flasks) were treated with ciprofloxacin and/or cephalexin. The samples were fixed with formaldehyde and incubated for 30 min with DAPI (2 μg/mL(Nonejuie et al., 2013)) and 0.1% TRITON to increase cell permeability. The fixed cells were put on agarose pads (2 % agar in water) and imaged on a Nikon Eclipse Ti epi-fluorescent microscope using a 100x oil objective (excitation 380-420 nm, emission >430 nm, exposure time 100 ms). Cell lengths, widths, and fluorescence intensity were extracted using the Fiji plug-in MicrobeJ (Ducret et al., 2016). For measuring the area of micro-colonies (Fig. S7) we used semi-automated ImageJ plugin JFilament (Smith et al., n.d.). After extracting the coordinates of the micro-colony contours from phase-contrast images, colony area was calculated as the area of the corresponding polygon (Lopez-Garrido et al., 2018; Ojkic et al., 2016).

### Computer simulations of the DNA replication model

The computer code used to simulate our model was written in Java. Each chromosome is represented as a one-dimensional lattice of *L*_0_ *=* 1000 sites. The ends of the lattice are either linked to each other (to represent a circular chromosome) or to another chromosome lattice at points corresponding to the current positions of the replication forks. Poisoned gyrases are identified by the index of the chromosome on which they sit, and their position (lattice site) within that chromosome. The simulation proceeds in discrete time steps (d*t* =*N*_bp_/(*L*_0_ *v*_f_), where *N*_bp_ =4,639,675 is the number of base pairs in the *E. coli* chromosome, and *v*_f_ = 30,000 bp/min is the fork speed. At each timestep, the position of each fork that can move (i.e. that is not blocked by a gyrase) is advanced by one lattice unit. Gyrases bind and detach with probabilities proportional to the corresponding rate times d*t*. Chromosomes are killed with probability *p*_kill_d*t* times the number of poisoned gyrases, and removed from the simulation. Chromosomes are separated when two forks reach the endpoints of the mother chromosome. A pair of new forks is added every *τ*_fork_ time units, where *τ*_fork_ is drawn from a normal distribution with mean 24 min and std. dev. 5 min. In simulations of the model with DNA damage occurring only at the forks, only stalled forks kill chromosomes (probability *p*_kill_d*t* per stalled fork).

All simulations were initiated with a single chromosome at *t* = 0 h, and stopped at *t* = 6 h (Figs. 3, S6, 7) or *t* = 5 h (Fig. 6). Between 1000 and 5000 independent runs were performed to obtain averaged curves. The step of CIP in Fig. 6 was simulated by running the simulation with *k*_+_ = 0 for *t* < 100 min, and switching to *k*_+_ > 0 corresponding to the desired CIP concentration for *t* > 100 min.

To fit the model to the experimental growth inhibition curves we systematically explored the space of parameters *p*_kill_ and *τ*_gyr_ (Fig. 3). The parameter *p*_kill_ was varied in the range 5.10^−5^ −10^−3^ min^-1^ for 11 data points, and *τ*_gyr_ was varied in the range 0 - 80 min in 5 min steps. For a given pair of values for *p*_kill_ and *τ*_gyr_ we simulated the model with different values of *k*_+_ and varied the scaling factor q to fit the experimentally obtained growth-inhibition curve by minimizing the sum of squares between the experimental and simulated inhibition curves. The best fit was obtained for *p*_kill_ = (7 +/- 2)10^−5^ min^-1^ and *τ*_gyr_= (25 +/- 5) min, *q* = (0.030+/- 0.005) ml ng^-1^ min^-1^ for the model with replication-independent killing, and *p*_kill_ = (2 +/- 1.5) 10^−5^ min^-1^ and *τ*_gyr_ = (30 +/- 5) min, *q* = (0.040 +/- 0.005) ml ng^-1^ min^-1^ for the model with replication-dependent killing (at the forks).

### Model for exponentially growing filaments (cephalexin)

To extract growth rates from the filament length distributions in Figs. 5 and 7, each cell was assigned an initial length *l*_0_ from the experimentally observed distribution (Fig. S4B), and a random growth rate *a* taken from a Gaussian distribution characterized by its mean and standard deviation (*α, σ*(*α*)). The new cell length after time *t* = 1h was calculated as *l* = *l*_0_ exp(at). A histogram of 642 000 predicted cell lengths was compared with the experimentally obtained cell length distribution for cephalexin-treated cells. The best match was obtained for *α* = 1.86 h^-1^ and *σ*(*α*) = 0.22 h^-1^ using the p-value from the Kolmogorov-Smirnov test as the goodness-of-fit measure. The best-fit mean growth rate was similar to the growth rate measured in the plate reader (1.7 h^-1^, Fig. 1A) indicating that cephalexin treated cells continued to elongate with the same rate for at least one hour in the presence of CIP. The spread of elongation rates given by *σ* (*α*) is similar to that observed for untreated cells (Taheri-Araghi et al., 2015; Wallden et al., 2016).

### Turbidostat

Our turbidostat device (Fig. S1C) encompasses 4 replicate cultures with a culture volume of approx. 26 mL. The growth medium used in all experiments was LB broth (Miller), and the *E. coli* strain used was AD30, to avoid biofilm formation. In the turbidostat, all cultures are connected to a bottle of LB medium and a bottle of LB + CIP (at 10x the desired CIP concentration in the culture) through a system of computer-controlled syringe pumps and valves. The optical density is measured every 20 s using custom-made photometers (separate for each bottle) to which 3-4 ml of each culture is aspirated and dispensed back into the culture using a syringe pump. When the optical density reaches OD=0.1 or after 30 min since the last dilution (whichever happens first), 25% of the culture is replaced with fresh medium to maintain exponential growth. An appropriate volume of CIP-containing LB medium is injected 2 hours after OD=0.1 is reached for the first time to achieve the required concentration (5-100 ng/ml) in the culture. Smaller volumes are injected in all subsequent dilution steps to maintain the prescribed concentration of CIP for the rest of the experiment. All cultures are kept in an incubator set to 37°C and are continuously stirred using magnetic stirrers and aerated with an air pump to keep dissolved oxygen (measured using Pyroscience FireStingO2) well above 50% of saturation concentration at 37°C (aerobic conditions).

## Acknowledgements

This work was supported by the European Research Council under consolidator grant 682237 EVOSTRUC. BW was supported by a Royal Society of Edinburgh Personal Research Fellowship. We thank Meriem El Karoui, Sebastian Jaramillo Rivera and Matthew Scott for helpful discussions, and Meriem El Karoui and Sebastian Jaramillo Rivera for supplying us with the *recA* mutant strain. This work has made use of resources provided by the Edinburgh Compute and Data Facility (ECDF; www.ecdf.ed.ac.uk).

## Supplementary Figures

**Figure S1.**
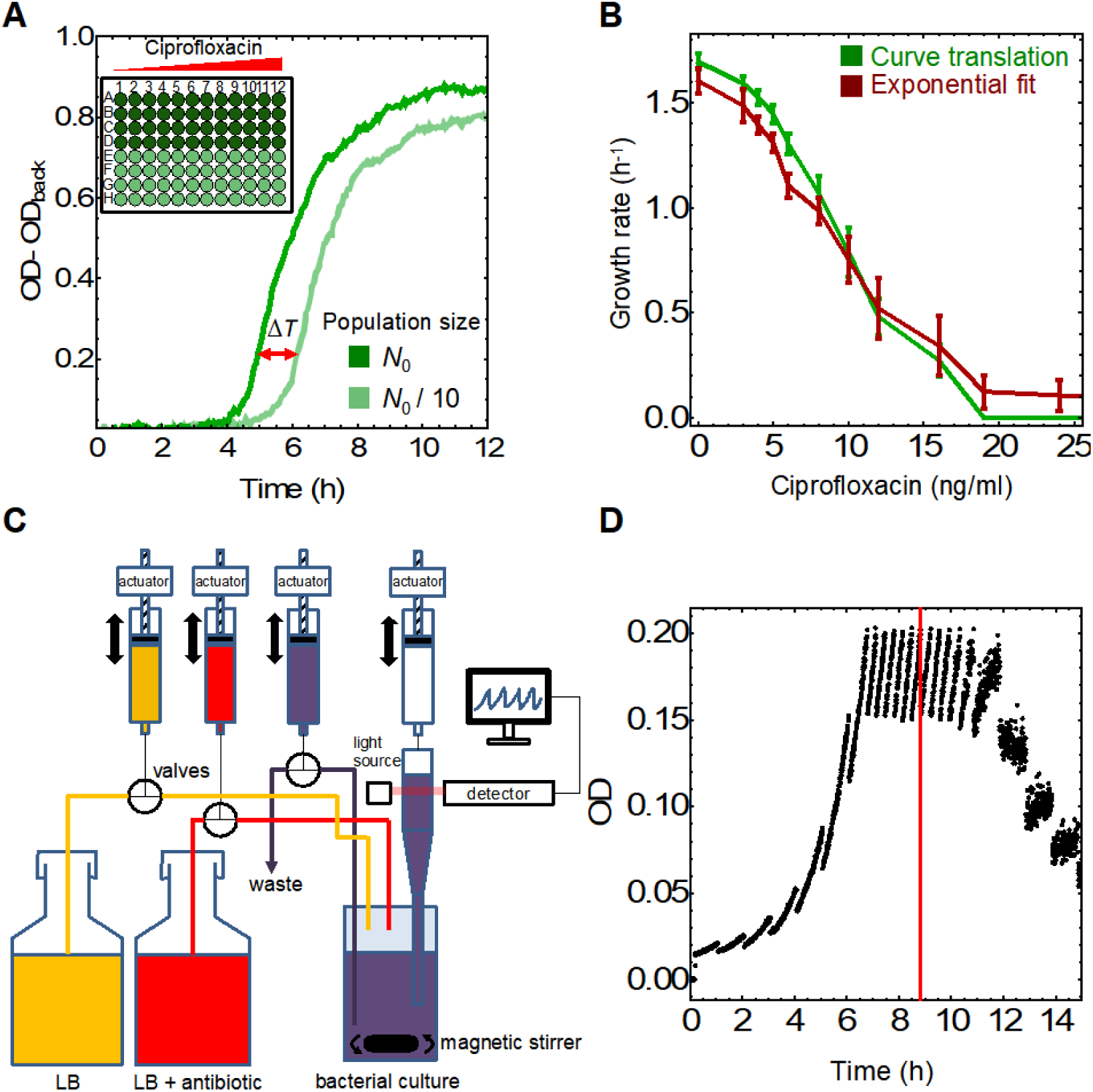
Growth rate measurements. (A) Background-corrected optical density OD_600nm_ vs time, measured in a plate reader for two initial population sizes (inocula) of *N*_0_ and *N*_0_/10 cells. The time delay (Δ*T*, red double arrow) is related to the growth rate via Eq. (1). (Inset) Microplate layout: columns = different concentrations, rows = different initial population sizes. (B) Growth-inhibition curve for ciprofloxacin-treated cells (MG1655). The minimum inhibitory concentration (MIC) is ∼20 ng/ml. Our time-shift method gives similar results to that of the standard exponential fitting method but it is more accurate (smaller error bars). Error bars are SEM. (C) Schematic drawing of the turbidostat. While only one bacterial culture is shown, the complete setup has four units that can be controlled independently. The pumps are syringe pumps (shared between the units), and computer-controlled valves control the flow to/from a particular unit. (D) Example data (OD versus time) from a single turbidostat experiment. The red line marks the time at which CIP was first added to the culture.

**Figure S2.**
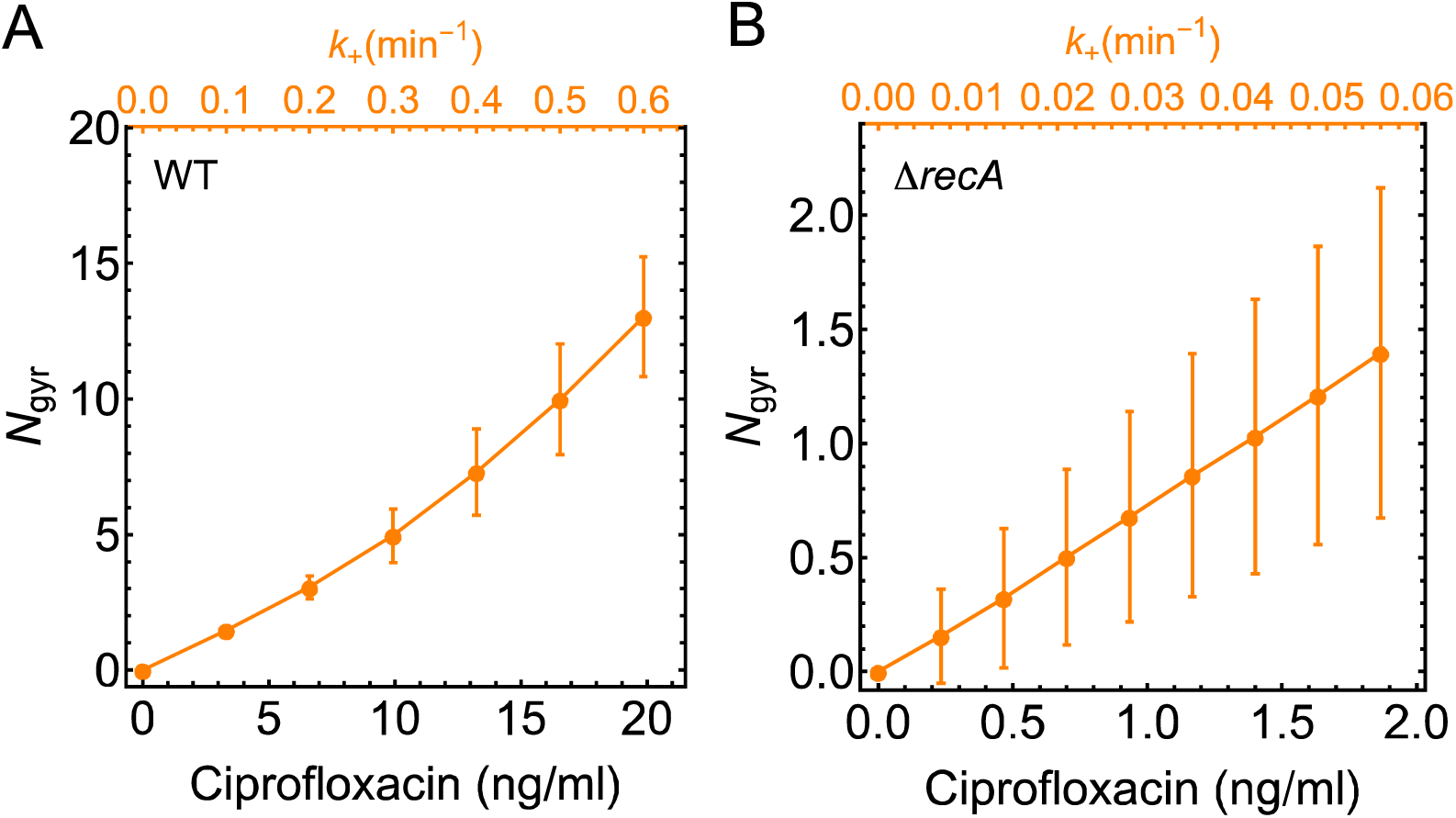
Number of poisoned gyrases predicted by the model. (A) For the best-fit parameters *p*_kill_ = 710^−5^ min^-1^ and *τ*_gyr_ = 25 min (Fig. 4), we calculated the average number of poisoned gyrases per chromosome length *<u>N</u>*_gyr_ (orange points, 1000 replicate simulations). (B) Same as in (A) but using the best-fit parameters for Δ*recA* cells (Fig. 7). According to the model, a single poisoned gyrase per chromosome is enough to cause complete DNA inhibition in cells lacking the recombination repair mechanism.

**Figure S3.**
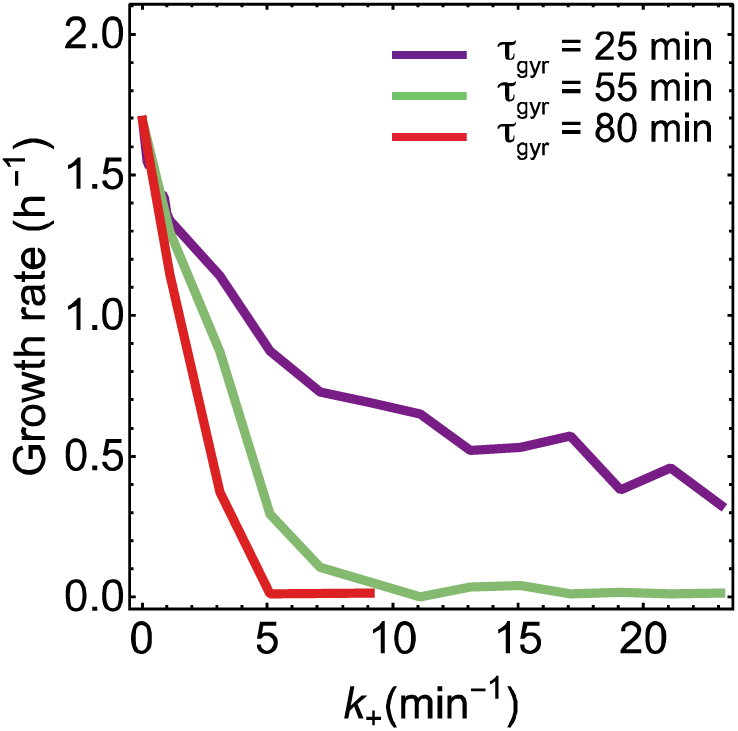
Simulation of the model when killing occurs only for the daughter chromosomes leaving the mother chromosomes intact. The predicted steep decrease in growth rate with CIP concentration is in sharp contrast to the quadratic shape of the experimental growth-inhibition curve from Fig. 1A.

**Figure S4.**
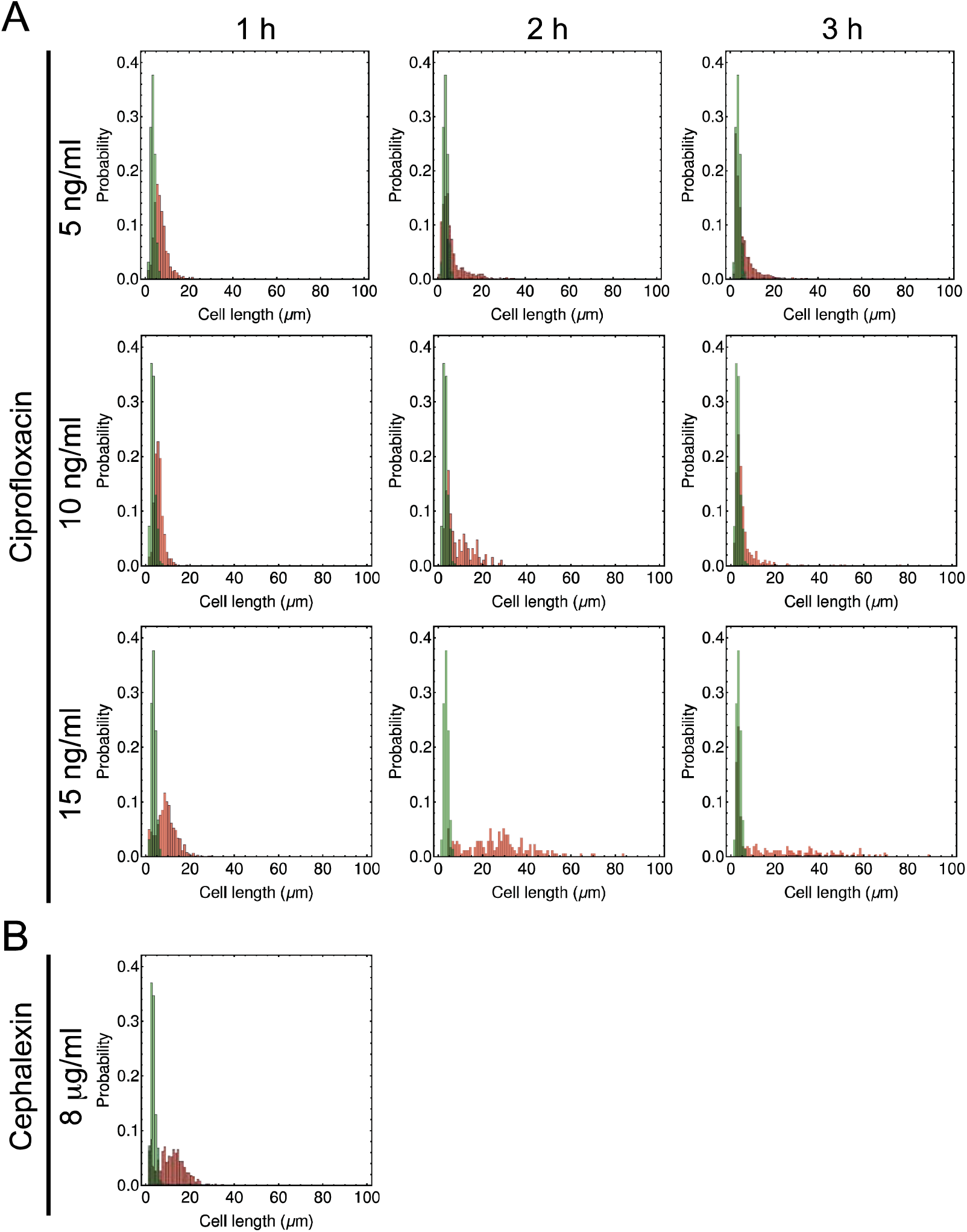
Cell length distributions for ciprofloxacin- and cephalexin-treated cells. The histograms show the cell length distributions before (green) and after antibiotic treatment (red). (A) When exposed to ciprofloxacin, cells form filaments that may bud from their end (Bos et al., 2015). Ciprofloxacin decreases the frequency of cell division; almost no cells bud or divide during first hour at the highest concentration used (15 ng/ml). (B) Cells exposed to 8 μg/ml (≈MIC) of cephalexin do not divide. The cell length distribution at *t* = 1 h is very similar to the distribution for 15 ng/ml of ciprofloxacin from panel A.

**Figure S5.**
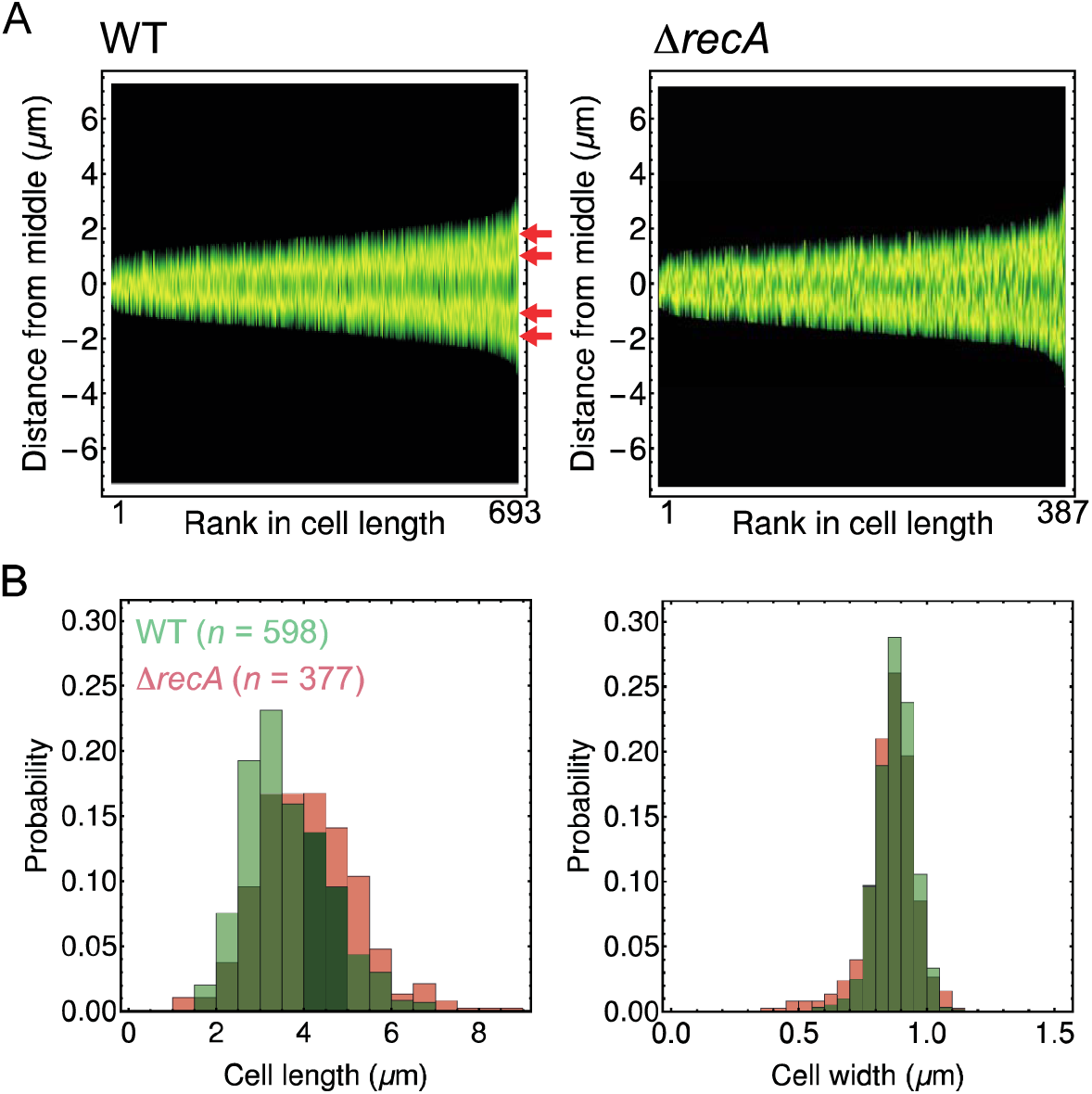
Chromosome organization in WT vs Δ*recA*. (A) Cells are ordered by length from shortest to longest along the *x*-axis, and fluorescence intensity (DAPI staining) is plotted along the *y*-axis. Isolated chromosomes (up to 4 in longest cells) can be identified in WT cells (red arrows), while Δ*recA* cells have much less organized chromosomes than WT cells. (B) The cell-length and cell-width distributions are very similar for both strains.

**Figure S6.**
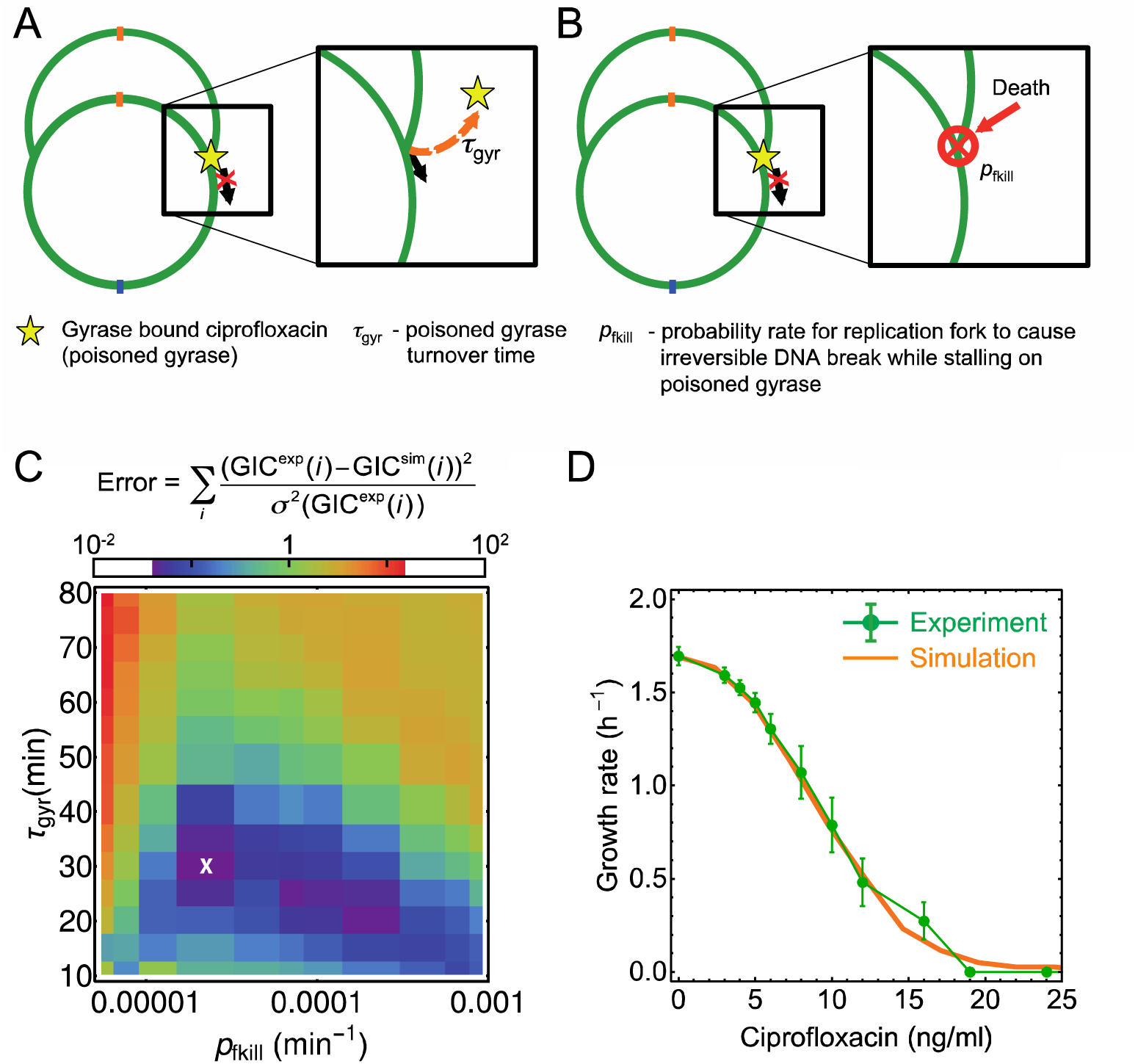
A model with DNA damage occurring at the stalled forks also reproduces the experimental growth-inhibition curve. (A) Schematic representation of the modified model, (c.f. Fig. 2). (B) Stalled replication forks cause irreversible DNA breaks with rate *p*_fkill_, leading to “death” of the chromosome. (C) Goodness-of-fit for a range of model parameters. The best-fit parameters *p*_kill_ = 2 10^−5^ min^-1^, *τ*_gyr_ = 30 min, and *q* = 0.04 ml ng^-1^ min^-1^ are marked with a white cross. (D) Experimental growth-inhibition curve (green) agrees well with the simulated curve (orange) for best-fit parameters. Errors are SEM (four replicates).

**Figure S7.**
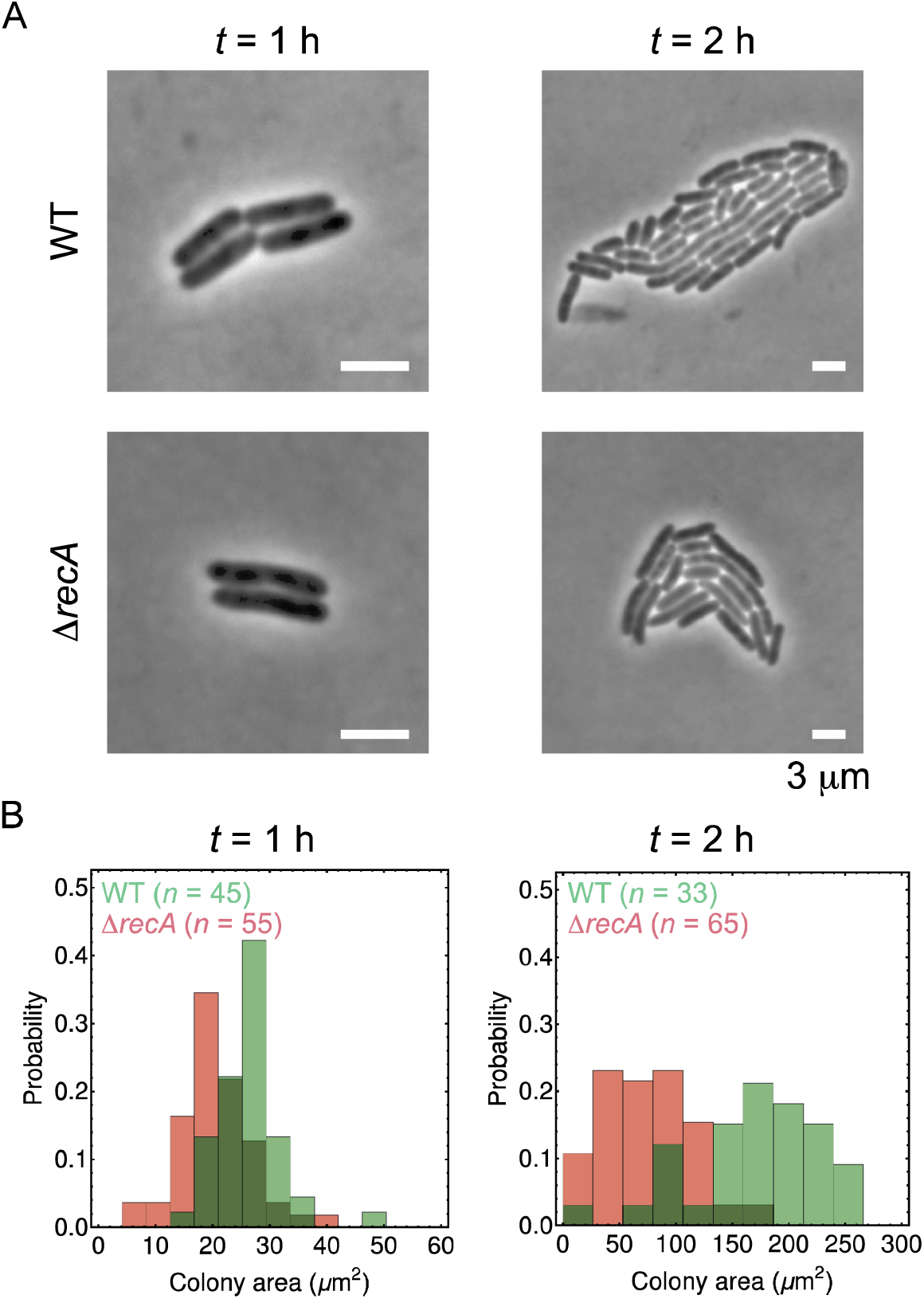
Colony size distribution for the WT (MG1655) and Δ*recA*. (A) Example colonies of WT and Δ*recA* cells imaged after 1 h and 2 h of growth starting from isolated cells deposited on LB-agarose pads. Scale bar = 3 μm. (B) Distribution of colony sizes. Colonies of Δ*recA* are smaller on average even though cells elongate with the same rate (Fig. 7B). By comparing the same colony at *t* = 1 and 2 h we concluded that some cells did not grow.

**Figure S8.**
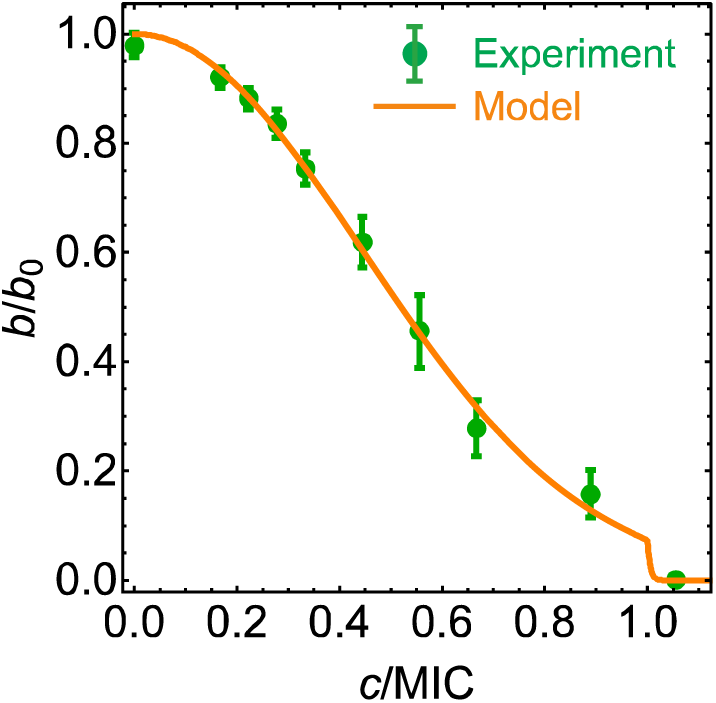
Alternative model (saturation of the repair mechanism). Experimental growth inhibition curve (green points) fitted with the model (orange line). Here *b*/*b*_0_ is the ratio of the growth rate at given CIP concentration *c* to the growth rate at *c*= 0. Although the inhibition curve is correctly reproduced, the model fails to reproduce the dynamic response as explained in the main text.

**Figure S9.**
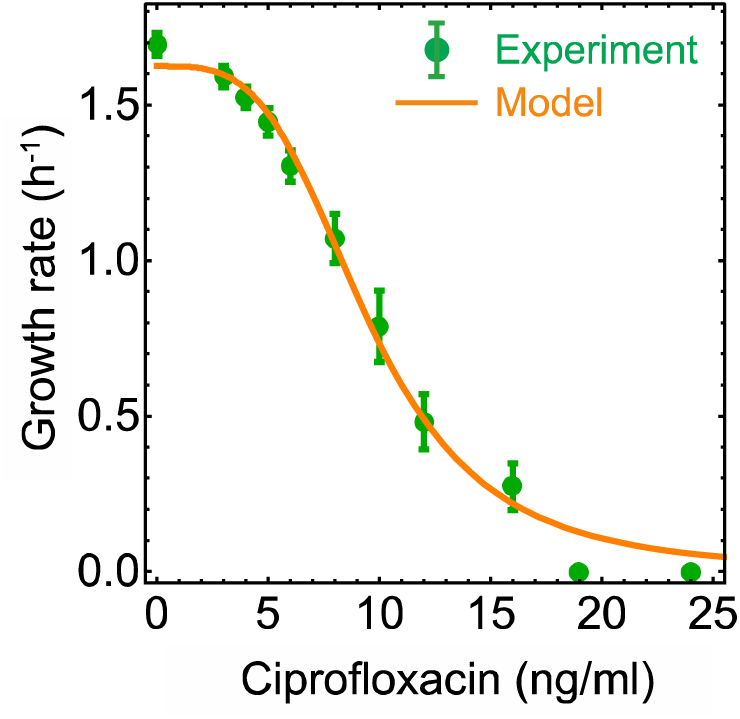
A Hill curve fitted to the experimental growth inhibition curve. The fitted Hill exponent is 4.4 ± 0.5.

**Figure S10.**
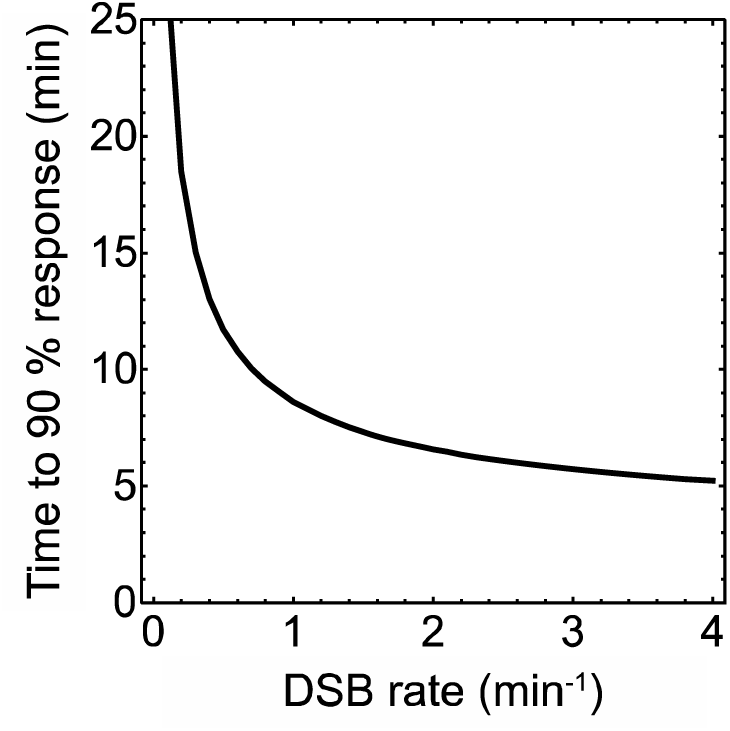
The SOS response is much faster than the experimentally observed growth response to CIP. The plot shows the time it takes the concentration of LexA (a protein involved in the SOS response) to reach its new steady state (less than 10% difference to the steady-state value) as a function of the rate with which DSBs are created. Based on model from (Belov et al., 2009) adapted as described in the main text.

